# Destabilization of F-actin by Mechanical Stress Deprivation or Tpm3.1 Inhibition Promotes a Pathological Phenotype in Tendon Cells

**DOI:** 10.1101/2022.02.15.480605

**Authors:** Kameron L. Inguito, Mandy M. Schofield, Arya D. Faghri, Ellen T. Bloom, Marissa Heino, Dawn M. Elliott, Justin Parreno

## Abstract

The actin cytoskeleton is a central mediator between mechanical force and cellular phenotype. In tendon, it is speculated that mechanical stress deprivation regulates gene expression by filamentous (F−) actin destabilization. However, the molecular mechanisms that stabilize tenocyte F-actin networks remain unclear. Tropomyosins (Tpms) are master regulators of F-actin networks. There are over 40 mammalian Tpm isoforms, with each isoform having the unique capability to stabilize F-actin sub-populations. Thus, the specific Tpm(s) expressed by a cell defines overall F-actin organization. Here, we investigated F-actin destabilization by stress deprivation of tendon and tested the hypothesis that stress fiber-associated Tpm(s) stabilize tenocyte F-actin to regulate cellular phenotype. Stress deprivation of mouse tail tendon fascicles downregulated tenocyte genes (collagen-I, tenascin-C, scleraxis, α-smooth muscle actin) and upregulated matrix metalloproteinase-3. Concomitant with mRNA modulation were increases in DNAse-I/Phallodin (G/F-actin) staining, confirming F-actin destabilization by tendon stress deprivation. To investigate the molecular regulation of F-actin stabilization, we first identified the Tpms expressed by mouse tendons. Tendon cells from different origins (tail, Achilles, plantaris) express three isoforms in common: Tpm1.6, 3.1, and 4.2. We examined the function of Tpm3.1 since we previously determined that it stabilizes F-actin stress fibers in lens epithelial cells. Tpm3.1 associated with F-actin stress fibers in native and primary tendon cells. Inhibition of Tpm3.1 depolymerized F-actin, leading to decreases in tenogenic expression, increases in chondrogenic expression, and enhancement of protease expression. These expression changes by Tpm3.1 inhibition are consistent with tendinosis progression. A further understanding of F-actin stability in musculoskeletal cells could lead to new therapeutic interventions to prevent alterations in cellular phenotype during disease progression.

## Introduction

The dysregulation of filamentous (F−) actin is associated with a wide range of pathologies in load-bearing mechanical tissues [1–10]. While the complete interactions between mechanical forces, F-actin, and pathology are not fully clear, the actin cytoskeleton may be a key mediator between mechanical force and cellular phenotype[1, 7, 11–14]. Tendon is an ideal tissue model to examine the mechanical regulation of cellular phenotype [15]. Mechanical loading of tendon is essential to maintain tissue homeostasis and tissue overloading is a cause of tendinosis [16–22]. While it may be speculated that tendon overload leads to mechanical over-stimulation of residing tendon cells, leading to alterations in cell phenotype (i.e., MMP production) [15, 23], an alternative hypothesis is that overload leads to a paradoxical mechanical under stimulation or stress deprivation of tendon cells [24, 25]. The hypothesized disruption of cell-matrix interactions during overload results in the failure of loads to be transmitted to cells. In support of this hypothesis, overloading of rat tail tendon resulted in cell-matrix disruptions as demonstrated using transmission electron microscopy by an increase in spacing between matrix and cells [26].

Mechanical loading/unloading of explant tendon tissue has been widely used to investigate the connection between cellular stress deprivation and tissue degeneration. Unloading or stress depriving tendons by culturing explants in the absence of mechanical load, recapitulates several aspects of tendinosis. Stress deprivation alters tendon cell phenotype causing a reduction in tendon molecule (such as collagen 1; Col1) expression and enhancement of matrix degradation molecule (such as matrix metalloproteinases; Mmps) expression as compared to tendons freshly isolated from a mechanically stressed *in vivo* environment [27–36]. However, stretching tendons while in culture prevents these tendinosis-like changes. The protective effects of stretch are abrogated if tendons are cultured in the presence of F-actin depolymerization agent, cytochalasin D, while being stretched [13, 37]. These findings have led to the speculation that stress deprivation regulates tendon cell phenotype via F-actin destabilization. This destabilization leads to depolymerization of F-actin into monomeric, globular (G-) actin which has been shown in other cell types to regulate gene expression [38–41]. However, there is limited data to support that stress deprivation destabilizes/depolymerizes F-actin. In the present study, we ask: is F-actin depolymerized during tendon stress deprivation? To answer this, we performed whole-mount confocal imaging followed by image quantification to compare freshly isolated stress-deprived tendons stained for F- and G-actin. We confirm that the acquisition of tendinosis-like molecular expression by stress deprivation correlates with F-actin destabilization.

Our finding of F-actin destabilization prompted us to examine the molecular regulation of F-actin network stability in tendon cells. We focused on the regulation of F-actin networks by tropomyosin (Tpm), the master regulator of F-actin that binds along and stabilizes F-actin [42]. The regulation of tenocyte F-actin by Tpms is largely unknown. In native and *in vitro* (primary cultured) tendon cells, F-actin organizes into stress fibers [14] containing alpha-smooth muscle actin (αsma), a highly contractile form of actin [11, 14, 43, 44]. Tpm associates with F-actin stress fibers in tendon cells [14]. However, the particular Tpm(s) expressed within tendon cells are not clear. Over forty isoforms of mammalian tropomyosin exist and each isoform has the unique capability to stabilize specific F-actin sub-populations. Thus, the particular Tpm(s) expressed within a cell defines overall F-actin organization.

In this study, we aim to determine which Tpm isoforms are expressed by tendon cells. Since tenocyte F-actin networks arrange into stress fibers, we focused on elucidating the function of stress fiber-associated Tpm isoforms. Previous studies using osteosarcoma (U2OS) cells have indicated at least six Tpms (1.6, 1.7, 2.1, 3.1, 3.2 and 4.2) are associated with stress fibers [45, 46]. In lens epithelial cells, we demonstrated that inhibition of just one Tpm isoform (Tpm3.1) prevents stress fiber formation resulting in downstream gene modulation [38]. Therefore, in this study we tested the hypothesis that stress fiber-associated Tpm(s) stabilize tenocyte F-actin and regulate cellular phenotype by modulating mRNA levels.

It is essential to unravel how cellular phenotype is determined by the F-actin network’s response to mechanical forces and to elucidate the molecules that determine/stabilize F-actin networks. Establishing the mechanisms involved in regulating cellular phenotype by F-actin in pathological processes, such as tendon overload, could lead to new therapeutic opportunities against disease.

## Methods

### Mice and tissue isolation

Wild-type mice, 8-10 weeks of age, in the C57BL/6J background were used for experiments. All procedures were conducted following approved animal protocols from the University of Delaware. Following euthanasia, plantaris, Achilles, and tail tendons were isolated as previously described [47–49]. To dissect plantaris and Achilles tendons, approximately 1 cm of the plantaris and Achilles complex were removed with the calcaneus intact. The tendons were separated by removing excess surrounding soft tissue before cutting off the calcaneal insertion [47]. Tail tendon fascicles were obtained from excised tails using forceps [48].

### Tendon explant stress deprivation culture

Tendon stress deprivation was performed by placing isolated tendon fascicles in culture vessels submerged in serum-free DMEM (GenClone, Genesee Scientific, San Diego, CA, USA) consisting of 1% antimycotic/antibiotic (MilliporeSigma, Burlington, MA, USA) [28, 33, 36, 50]. Tendons were maintained at 37°C and 5% CO_2_ and were harvested at set time points, either 1- or 2-days post-isolation. For these studies, tail tendon fascicles were distributed in a non-biased fashion to experimental conditions, with freshly isolated fascicles serving as controls.

### Tail tenocyte isolation and Tpm3.1 inhibition by TR100 treatment

To obtain tendon cells, tail tendon fascicles were isolated and immersed in 0.2% Collagenase A (MilliporeSigma) and maintained at 37°C. Following overnight incubation, digests were strained through 100μm filters and then centrifuged at 600g for 6 minutes. Tenocyte pellets were resuspended in fresh DMEM consisting of 10% fetal bovine serum (FBS; GenClone) and 1% antimycotic/antibiotic. For experiments, cells were counted and then seeded on culture vessels at a density between 5 to10 × 10^3^cells/cm^2^. Every two days spent media was replaced with fresh media. Once tenocyte cultures reached ∼70% confluency, cells were serum starved in DMEM consisting of 0.5%FBS and 0.1% antimycotic/antibiotic with or without Tpm3.1 inhibitor (TR100; 1-10μM; MilliporeSigma). Tendon cells were harvested 1-or 2-days after drug treatment.

### Light microscopy and cell area/circularity quantification

Following 24 hours of treatment, cells were imaged using a microscopy camera (Swiftcam Technologies, Hong Kong) attached to an Axiovert 25 inverted phase-contrast microscope (Zeiss, Jena, Germany). The boundaries of individual cells were manually traced on images using FIJI software. Cell area and circularity were calculated using FIJI software algorithms as previously performed [39]. Circularity was defined as C = 4π(A/P^2^) where, P is the perimeter, and A is the cell area. A circularity value of 0 indicates an elongated elipse, whereas a value of 1 indicates a perfect circle.

### Cell viability assay

We used a propidium iodide exclusion assay to quantify the proportion of live cells following TR100 treatment. Cells on culture vessels were placed in 0.05% trypsin at 37°C. After 5 minutes, cells were centrifuged, and pellets were then resuspended in PBS containing 0.5μg/mL propidium iodide. Propidium iodide is impermeable to live cells but incorporates into dead cells with compromised cell membranes. A Tali Image-based Cytometer (ThermoFisher Scientific; Waltham, MA) was used to calculate the portion of cells in suspension that incorporated propidium iodide. Cells exposed to 70% ethanol served as dead controls.

For visualization of cell viability, cells on culture vessels were stained with 1μM Calcein AM (live cell stain) and 1μM propidium iodide in PBS at 37°C. After 10 minutes, cells were rinsed with PBS and visualized using a Zeiss Axio Observer 7 microscope. Cells fixed in 4% paraformaldehyde (before staining) were used as dead cell control.

### RNA isolation, gene expression, and Tpm sequencing

For RNA processing, tendon tissue or cultured tendon cells were immersed in TRIzol (Sigma-Aldrich, Burlington, MA) and stored frozen until further use. To extract RNA from tissues, tissues were manually grinded in TRIzol using a Pellet Pestle. RNA was separated via phase separation with chloroform, followed by selective recovery of total RNA using an RNA spin column clean-up kit (RNA Clean & Concentrator-5; Zymo, Irvine, CA). RNA was extracted from cultured cells by phase separation with chloroform followed by RNA precipitation in isopropanol as previously described [38]. Reverse transcription of RNA to cDNA was performed using UltraScript 2.0 cDNA Synthesis Kit (PCR Biosystems, Wayne, PA).

Semi-quantitative RT-PCR was performed using PCRBIO Taq Mix Red solution (PCR Biosystems; London, UK) on equivalent amounts of cDNA using previously validated Tpm primers [51]. PCR products were run on a 2% Agarose gel. Gel bands were excised, melted, and DNA was purified using GenCatch Advance Gel Extraction Kit (Epoch Life Sciences; Missouri City, TX, USA). Purified DNA was sequenced using Sanger sequencing (Genewiz, South Plainfield, NJ, USA).

Relative real-time RT-PCR was performed using qPCRBio Sygreen Blue Mix (PCR Biosystems, Wayne, PA) according to manufacturer’s directions. Each reaction contained 20ng of cDNA in a 10μL volume using primers that are listed in Table 1. Real-time PCR was conducted on a Cielo 3 PCR machine (Azure; Houston, TX, USA).

**Table 1.**
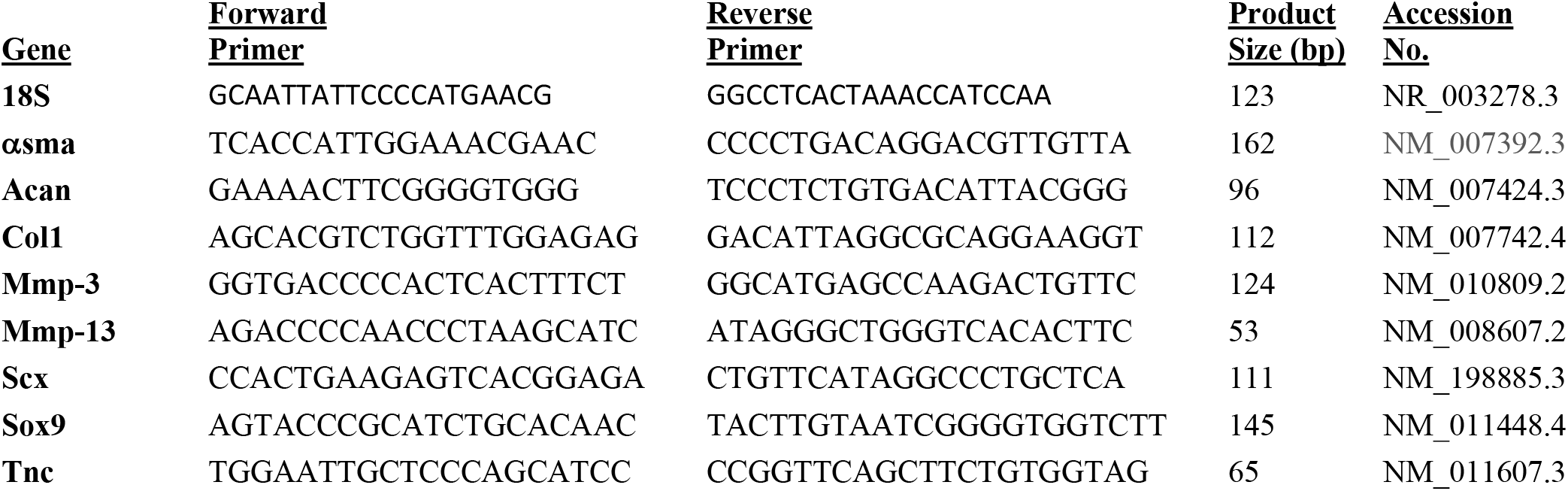
List of Real-time RT-PCR Primers

### Wes capillary electrophoresis

Total protein was extracted as previously described [38] with slight modifications. Briefly, protein was extracted in 1x Radioimmunoprecipitation (RIPA) lysis buffer (Millipore Sigma) and then quantified using a bicinchoninic acid protein assay (BCA) (Prometheus Protein Biology Products; Genesee Scientific, San Diego, CA, USA). Specific protein levels were determined using a WES Capillary-Based Electrophoresis assay with a 12-230 kDa separation module kit following the manufacturer instructions (Protein Simple, San Jose, CA, USA) with an equal volume of control and experimental protein. The antibodies used for experiments are rabbit anti-pan-Actin (1:100; #4968, Cell Signaling), mouse anti-αSMA (1:50; ab7817, Abcam), and rabbit anti-MMP-3 (1:10; EP1186Y, Abcam). Secondary antibodies used were anti-rabbit or anti-mouse HRP-conjugate (Protein Simple). Specific protein levels were normalized to total protein. Total protein values were determined using a Chemiluminescent WES Simple Western Size-Based Total Protein Assay (Protein Simple).

### Immunostaining of native tendon

Whole tendons were placed in 4% paraformaldehyde in phosphate-buffered saline (PBS; GenClone) at 4°C for 2 hours. Fixed tendons were then washed three times in PBS and incubated in permeabilization/blocking buffer (PBS containing 0.3%Triton, 0.3% bovine serum albumin, and 3% goat serum) at room temperature. After 2 hours, tendons were transferred into a primary antibody solution containing 1:100 mouse anti-Tpm3.1 antibody (2G10.2; Millipore Sigma) in permeabilization/blocking solution. After overnight incubation at 4°C, tendons were washed six times in PBS and placed in secondary antibody solution containing fluorescent CF488 anti-mouse secondary antibody (1:200; Biotium, Fremont, CA), Hoecsht 33342 (1:500; Biotium), and rhodamine-phalloidin (1:20; Biotium) in permeabilization/blocking buffer at room temperature for 2 hours in the dark. Tissues were then washed six times in PBS, placed on a glass slide, and mounted using ProLong gold anti-fade reagent (Thermo Fisher Scientific).

To stain for G/F-actin, fixed tendons were placed in permeabilization/blocking buffer at room temperature. After 2 hours, tendons were transferred into permeabilization/blocking buffer containing Hoecsht 33342 (1:500), Alexa Fluor DNAse-I (Thermo Fisher Scientific) and rhodamine-phalloidin (1:20).

### Immunostaining of cultured cells

Tendon cells cultured on glass-bottomed dishes (World Precision Instruments, Sarasota, FL, USA) were fixed with 4% paraformaldehyde in PBS at room temperature. After 10 minutes, fixed cells were washed in PBS and kept at 4°C. Tendon cells were immunostained by first incubating in permeabilization/blocking buffer at room temperature for 30 minutes. Next, cells were incubated with mouse anti-TPM3.1 antibody (1:200; 2G10.2, MilliporeSigma) or rabbit anti-SCX antibody (1:100; ab58655, Abcam) diluted into permeabilization/blocking solution. After overnight incubation at 4°C, cells were washed six times in PBS and placed in secondary antibody solution containing CF488 anti-mouse secondary antibody (1:200), Hoechst 33342 (1:500) to stain nuclei, and rhodamine-phalloidin (1:50) to stain F-actin, in permeabilization/blocking buffer at room temperature for 1 hour in the dark. Cells were then washed six times in PBS and mounted using ProLong gold anti-fade reagent (Thermo Fisher Scientific).

### Confocal fluorescence microscopy

Stained tissues and cells were imaged using a Zeiss LSM880 laser-scanning confocal fluorescence microscope (Zeiss) using a 20x 0.8NA objective or a 63x 1.4NA oil objective. Z-stack images were acquired using 0.5μm (20x objective) or 0.3μm (63x objective) step size. Images were processed using Zen software from Zeiss.

G/F-actin in whole-mount confocal images was quantified using FIJI software. Briefly, single optical slice images from the mid-region of full z-stack tendon images were analyzed. A rectangular region of interest (ROI) was selected at the center of the images and channels (red/green/blue) were then separated. The fluorescent intensities within the ROI for each channel were measured. The fluorescent intensity in the green (DNAse-I) channels were divided by the fluorescent intensity in the red (Phalloidin) channels to obtain DNAse-I/Phalloidin (G/F-actin) ratios. Finally, the G/F-actin ratios obtained from each ROI were normalized to the average G/F-actin control values (control averages were set to 1). Of note, tendons from control and stress-deprived groups were stained using identical conditions simultaneously, images were acquired using the same settings on the same day, and ROIs were equivalent in size.

### Statistical analysis

Each experiment was replicated at least three times on separate occasions. Data from individual experiments were expressed as a percentage of the control average, which was set to 100. Graphpad Prism 9 (San Diego, CA, USA) was used for statistical analysis. Outliers in pooled data points were identified using the ROUT method with a maximum desired false discovery rate (Q) set at 1% [52]. Differences between two groups of data were detected using unpaired t-tests. Differences between several experimental groups against a control were detected using analysis of variance followed by Dunnett post hoc tests.

## Results

### Tendon stress deprivation results in tendinosis-like gene changes

Tendinosis progression is characterized by decrease in tenogenic gene expression, an upregulation of degradative genes, and ectopic expression of cartilage genes [27–36, 53–55]. Tendon stress deprivation, *in vitro*, has been shown to recapitulate several aspects of tendinosis gene modulation. We evaluated the effect of stress deprivation on tail tenocyte gene expression by culturing mouse tail tendon fascicles in the absence of stretch for up to 2 days.

Stress deprivation decreases tenogenic mRNA levels (Figure 1A). We observe a decrease in Col1, the predominant collagen expressed in tendon [54, 56, 57]; and a decrease in tenascin C (Tnc), a mechanosensitive matricellular glycoprotein present in tendon matrix that assists in the structural arrangement of collagen fibers as well as provides for tissue elasticity [12, 56, 58]. In addition, we observe a reduction in Scleraxis (Scx), a transcription factor that drives the tenogenic expression, such as the expression of collagen-I [59, 60]. Finally, we detect a decrease in alpha-smooth muscle actin (αsma), an actin isoform present in tendon cells, which is thought to be necessary for recovery of tendon cells following stretch [54].

**Figure 1.**
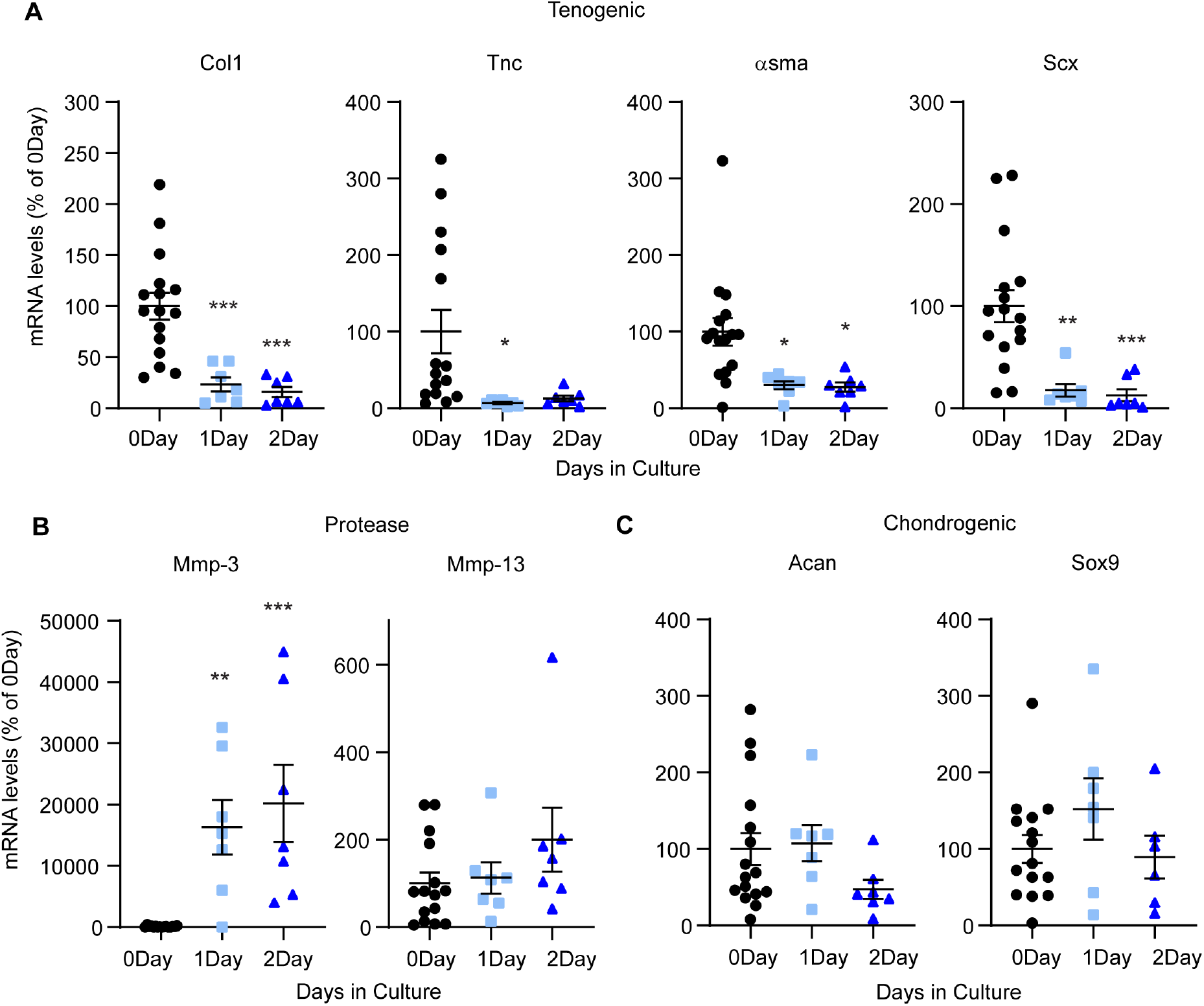
Stress deprivation recapitulates aspects of tendinosis gene expression. Relative real-time PCR of tendons in 1 or 2 day stress deprivation culture as compared to freshly isolated (0 day) tendons demonstrating (A) a reduction in tenogenic marker mRNA levels, (B) an increase in MMP-3 mRNA levels, and (C) no change in chondrogenic mRNA levels. Specific mRNA levels were normalized to Gapdh and normalized values are expressed as a percent of freshly isolated (0 day) controls. *, p < 0.05; **, p < 0.01; ***, p < 0.001

Stress deprivation enhances the expression of specific degradative molecules (Figure 1B). Mmp-3 and Mmp-13 have been implicated in tendinosis and are upregulated during stress deprivation [13, 23, 30, 35, 57, 61]. We determine that Mmp-3 is upregulated 163.1-fold by day 1 and 201.9-fold by day 2 (Figure 1B). Unlike previous studies [30], we did not observe an increase in Mmp-13 mRNA levels at 1- or 2-days post culture.

An increase in chondrogenic expression is a feature of tendinosis [55] but is not recapitulated by tendon stress deprivation. We observed no significant effect on chondrogenic (Acan or Sox9; Figure 1C) mRNA levels up to 2 days of stress deprivation (Figure 1C). Thus, *in vitro* stress deprivation of tail tendons recapitulates certain, but not all, aspects of tendinosis.

### Stress deprivation promotes cell rounding and destabilizes tenocyte F-actin

We found that tendon stress deprivation alters tenocyte morphology and F-actin depolymerization [6, 11, 13]. To determine if F-actin depolymerization occurs in native tendons, we performed whole-mount imaging of tendons stained with DNAse-I and Phalloidin to visualize G-and F-actin, respectively. Tendon cells from freshly isolated tendons are elongated and have parallel F-actin networks regarded as stress fibers throughout the tissue [14] (Figure 2A, B). Stress deprivation results in rounded tendon cells. We also observe repression of stress fiber organization. Fluorescent image quantification reveals that stress deprivation elevates G/F-actin staining after one day compared to freshly isolated controls (Figure 2C). By two days of culture, despite differences in F-actin network organization, G/F-actin status is similar to controls (data not shown). Our findings confirm that stress deprivation destabilizes F-actin.

**Figure 2.**
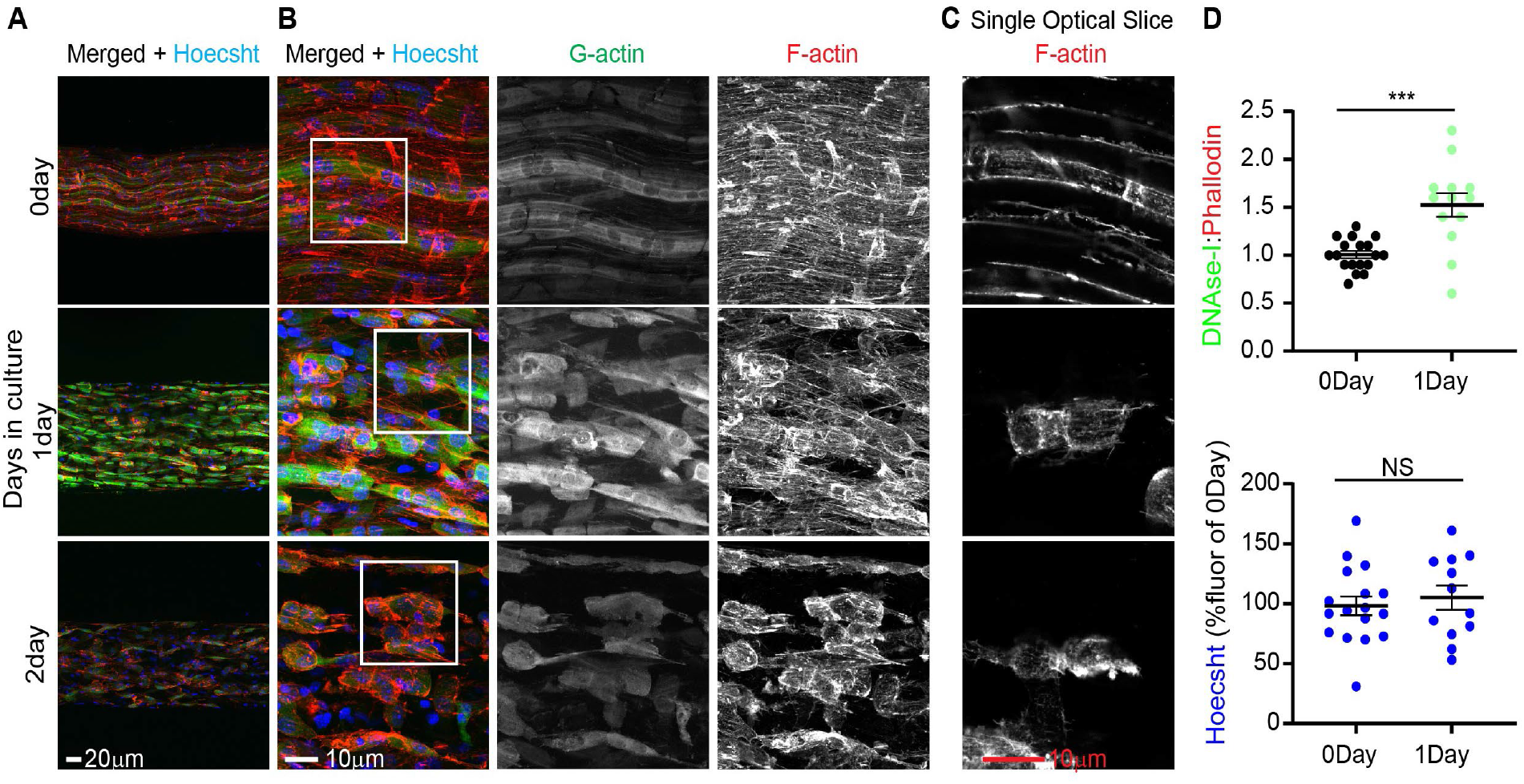
Stress deprivation alters tendon cell morphology and destabilizes F-actin. (A-B) Maximum intensity projections images of tendons demonstrating an increase in G/F-actin staining after 1 day of culture taken with either a 20x (panel A) or 63x (panel B) objective. In panel B, gray scale images represent individual channels for G- and F-actin. (C) Single optical slice of boxed region in panel B, demonstrates that cell morphology and F-actin are altered by stress deprivation. (D) Dot plots of DNAseI:Phalloidin to demonstrate an increase in G/F-actin. Average DNAseI:Phallodin for controls were set to 1.0 and 1 day stress deprived values were scaled accordingly. Nuclei staining intensity using Hoechst was calculated to demonstrate there were no significant differences in staining between control and stress deprived tendons. Average fluorescent intensity for controls were set to 100% and stress deprived tendon intensities were expressed as a relative percentage to controls. ***, p < 0.001

These results support the idea that F-actin destabilization regulates tendinosis gene changes. Tpms are known master regulators of F-actin, therefore we next focused on characterizing the role of Tpms in tendon cells.

### Tpm Expression in Native Tendon cells

Tpm isoforms have the unique ability to stabilize specific F-actin networks [42, 46, 62, 63]. Therefore, we first sought to determine which Tpm isoforms are expressed by tendon cells. Tpm expression is variable and dependent on tendon type. To determine specific Tpm isoform expression within various tendon types (tail, Achilles, and plantaris) tendons, we used semi-quantitative RT-PCR analysis, followed by Sanger Sequencing of cDNA products. Collectively, in total, we found that tendons express 9 Tpm isoforms: Tpm1.1, 1.6, 1.8, 2.1, 2.2, 3.1, 3.5, 3.12, and 4.2. However, the specific Tpms expressed are dependent on tendon type. Only three of the nine Tpm isoforms are consistently expressed in all three tendon types tested: Tpm1.6, 3.1, and 4.2.

In this study we chose to focus on the regulation of F-actin networks by Tpm3.1, as we previously have shown it to regulate stress fibers in lens epithelial cells [38]. Furthermore, there are established inhibitors toward Tpm3.1, including TR100 [64, 65]. To confirm Tpm 3.1 protein expression and association with F-actin, we performed whole-mount immunostaining of the tail fascicle using antibody 2G10.2 (Table 2) which recognizes the 9D exon of Tpm3.1. Our findings indicate Tpm 3.1 expression within native tail tendon cells. Furthermore, line scan analysis shows overlap in signal between Tpm3.1 and phalloidin staining, suggesting that Tpm3.1 associates with native F-actin stress fibers (Figure 3C).

**Figure 3.**
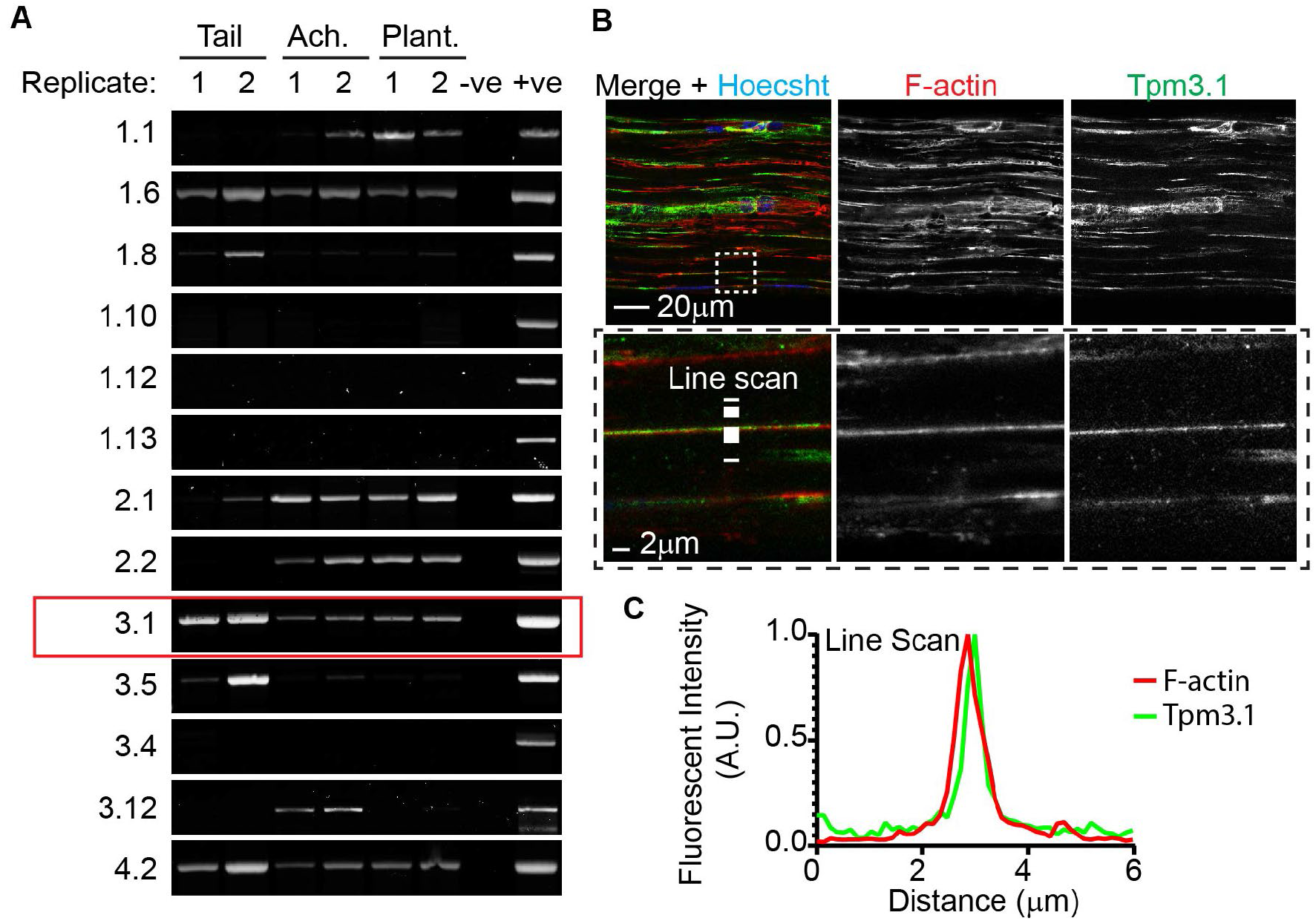
Tpm isoform expression is dependent on tendon type, but all three types examined express Tpm3.1. (A) Semi-quantitative PCR for Tpms in native tail, Achilles (Ach.), and plantaris (Plant.) tendons. (B, top panels) Whole mount immunostaining shows distribution of Tpm3.1 throughout tail tendon. (B, bottom panels) High magnification images of dashed area shows association of Tpm3.1 along F-actin. (C) Line scan analysis showing co-localization of Tpm3.1 with F-actin.

### Inhibition of Tpm3.1 with TR100 alters tenocyte morphology

To determine Tpm3.1 function in tendon cells, we used primary tendon cells isolated from tail tendon fascicles. Primary tendon cells retain stress fiber organization of F-actin [14]. Furthermore, cultured cells devoid of matrix are more amenable to biochemical assays (i.e., Triton-based cellular fractionation) than native tissues.

We exposed cultured tendon cells to TR100, a small molecule inhibitor that prevents end-to-end assembly of Tpm and reduces Tpm binding along F-actin [64, 66]. Live/dead staining (Figure 4A) and propidium iodide exclusion assay (Figure 4B) demonstrate that treating tendon cells with ≤5uM of TR100 retains cell viability. To determine the effect of Tpm3.1 inhibition on cell morphology, we performed light microscopy followed by image quantification (Figure 4C). Both 2μM and 5μM TR100 decrease overall cell area (Figure 4D). Interestingly, while 2μM TR100 induces tenocyte elongation (decrease in circularity), exposure to 5μM TR100 causes cell rounding (increase in circularity).

**Figure 4.**
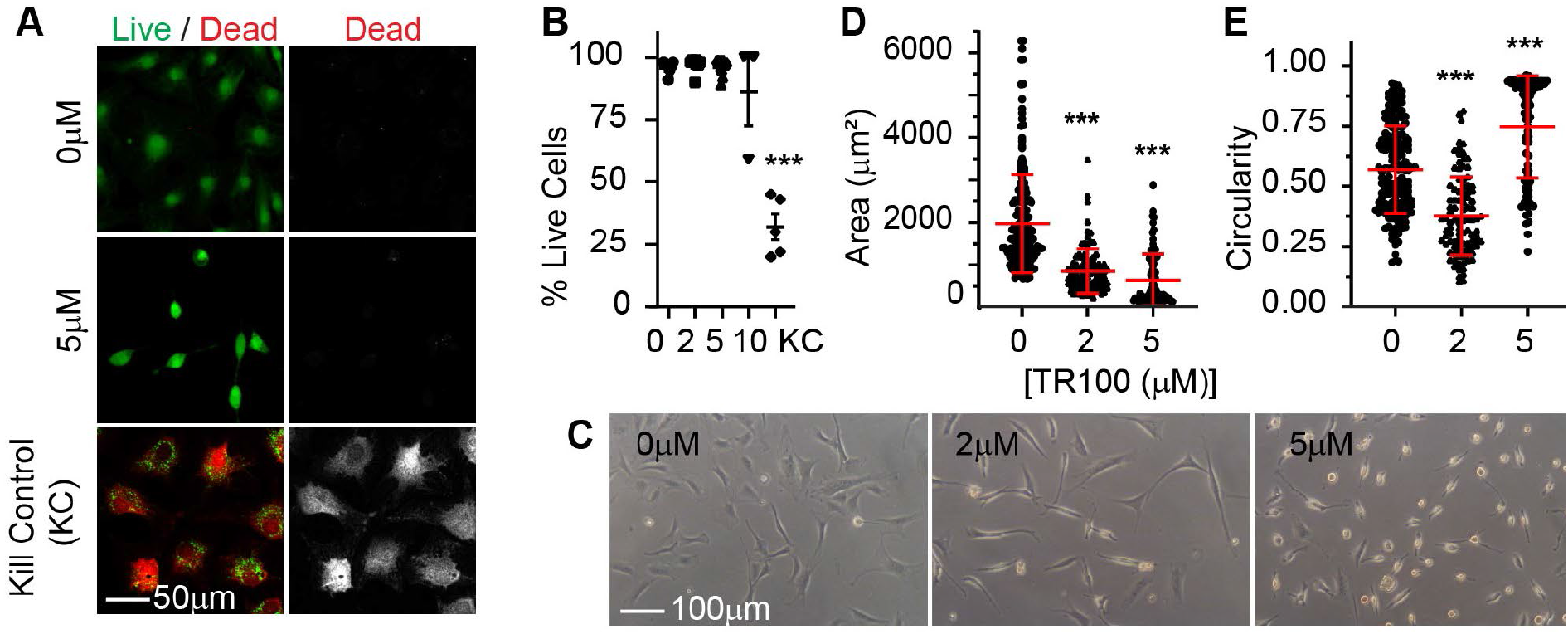
Exposure of cells to TR100 alters tenocyte shape and size. (A) Tendon cells are viable following 1 day treatments of TR100 up to 5mM as seen by positive staining for live cell stain (Calcein AM; green) and negative staining for dead cell stain (Propidium Iodide; red). Cells fixed in 70%EtOH for 15 minutes served as dead controls (Kill control; KC). (B) Dot-plot showing the percentage of cells that are alive following treatments as measured by propidium iodide exclusion counts. (C) Light microscopy images of cells treated with TR100 and corresponding dot-plots demonstrating changes to tenocyte (D) cell area and (E) circularity. ***, p < 0.001

### Tpm3.1 stabilizes Tenocyte F-actin

We sought to determine if Tpm3.1 stabilizes tenocyte F-actin. Confocal microscopy demonstrates that stress fibers are present in cells treated with 2μM TR100 (Figure 5A). However, there appears to be a slight reduction in Tpm3.1 staining along F-actin stress fibers compared to control (Figure 5B). Treatment with 5μM TR100 represses F-actin stress fibers and results in cortically arranged Tpm3.1 and F-actin in a subpopulation of cells.

**Figure 5.**
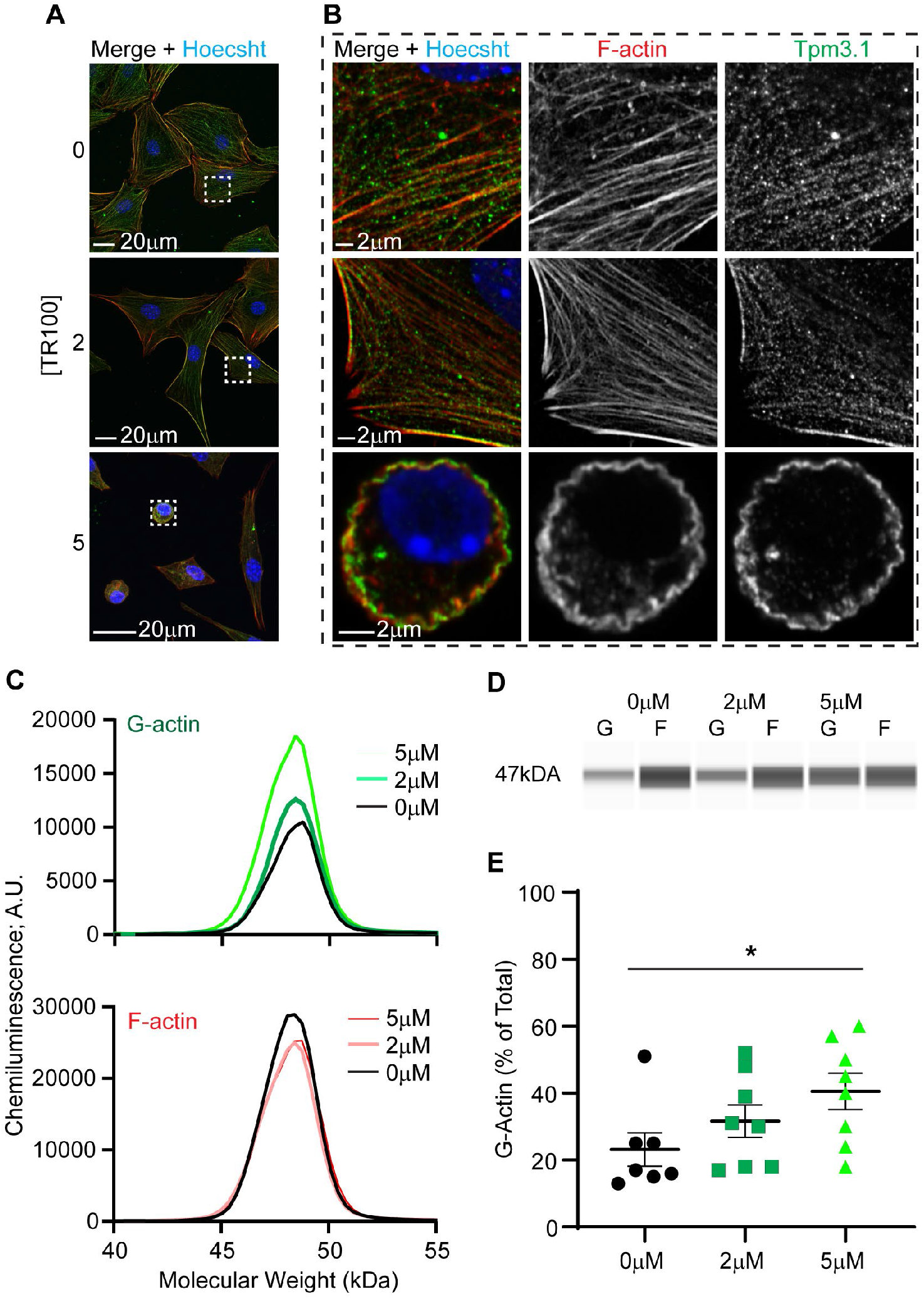
Tpm3.1 inhibition destabilizes F-actin. (A) Maximum intensity projection images of TR100 treated tendon cells, demonstrating that TR100 represses F-actin stress fiber organization. Images taken with a 20x objective. (B) Single optical slice images of boxed regions, demonstrating repression of F-actin stress fibers and acquisition of cortical organization of Tpm3.1 and F-actin (5μM TR100). Images taken with a 63x objective. (C) Capillary western blot (WES) immunoassay electropherogram of Triton-fractionated samples. The triton-soluble fraction is indicative of G-actin (top panel) and the triton-insoluble fraction is indicative of F-actin. (D) Virtual gel showing actin bands at approximately 47kDA. (E) Corresponding dot-plots showing the proportion of G-actin within cells increases by treatment with 5μM TR100. *, p < 0.05

Next, we evaluated G/F-actin proportion in TR100 treated cells using Triton-fractionation followed by WES-capillary electrophoresis (Figure 5C-E). Treatment with TR100 increases the amount of actin present in soluble portions indicative of elevated G-actin (Figure 5D, E). On average, there is a significant increase in the proportion of G/F-actin by treatment with 5μM of TR100. While 2μM of TR100 appears to result in a slight increase in the proportion of G/F-actin, this was not statistically significant (p = 0.42).

### TR100 treatment results in tendinosis-like gene changes

Our findings in Figure 5 reveal that Tpm3.1 regulates F-actin stability in tendon cells. The findings that F-actin destabilization regulates tendinosis-like gene changes, leads to us next testing the hypothesis that Tpm3.1 inhibition will induce downregulation of tenogenic genes and upregulation of chondrogenic and protease expression.

We determined that destabilization of F-actin by TR100 treatment decreases tenogenic mRNA levels. This was apparent at both 2μM and 5μM treatments. With 5μM treatment of TR100, there is a 21.2-, 1.4-, 29.2-, 2.5-fold decrease in Col1, Tnc, αsma, and Scx respectively. Treatment with 5μM of TR100 also upregulates chondrogenic mRNA levels resulting in a 3.4- and 9.2-fold increase in Acan and Sox9, respectively. Lastly, exposure of cells to 5μM TR100 upregulates protease expression resulting in a 18.1 and 374.3-fold increase in Mmp-3 and Mmp-13, respectively. We found that only the 5μM treatment, but not 2μM treatment, upregulates chondrogenic and protease levels.

We examined modulation of protein expression by 5μM TR100 treatment on select molecules using immunostaining followed by quantitative image analysis (SCX) or via capillary electrophoresis (αSMA and MMP-3). TR100 decreases the total nuclear fluorescent staining for SCX in tendon cells (Figure 7A, B) as well as the proportion of nuclear/cytosplasmic SCX (Figure 7A, C). Additionally, TR100 decreases αSMA protein levels by 2.0-fold (Figures 7D-F). Lastly, TR100 increases MMP-3 protein levels by 14.3-fold (Figure 7D-F). Modulation of SCX, aSMA, and MMP-3 is consistent between mRNA and protein levels.

## Discussion

This study provides novel insight into the regulation of pathological cellular phenotype by F-actin destabilization. We determined tendinosis-like gene changes caused by stress deprivation coincides with F-actin destabilization. This result led us to investigate the molecular regulation of F-actin stability. We revealed that tendons express several Tpm isoforms, including Tpm3.1, a stress fiber associated Tpm [38, 45, 46]. We found that Tpm3.1 associates with F-actin networks in tendon cells and regulates tenocyte cellular phenotype. Tpm3.1 inhibition destabilizes tenocyte F-actin stress fibers causing a robust tendinosis-like gene response.

Our data supports the speculation that stress deprivation regulates cellular phenotype by destabilization of F-actin [37]. Previously it was shown that tendon stress deprivation increases Mmp mRNA levels [28, 37]. It was also demonstrated that stretching *in vitro* cultured tendons can repress Mmp upregulation [28, 37]. Mechanical stretch appears to regulate Mmp expression via actin-based mechanisms as treating stretched tendons with cytochalasin D (an actin depolymerization agent) abrogates the protective effects of stretch [37]. This led to the speculation that stress deprivation regulates Mmp expression via actin depolymerization. However, evidence for F-actin depolymerization in stress deprivation was still required. In the current study, we provide empirical evidence that stress deprivation destabilizes F-actin as demonstrated by an increase in G/F-actin staining (Figure 2). We show this increase in G/F-actin coincides with an increase in protease expression (Mmp-3) (Figure 1). This finding is consistent with our previous findings in bone cells, where stress deprivation of osteoblast-like MG-63 cells in collagen gels enhances Mmp-3 gene expression [61]. Unlike other previous studies, Mmp-13 was not found to be upregulated, although there was a trend toward an increase in Mmp-13 expression by day 2 (p = 0.17). Previous studies examined Mmp expression at later time points (i.e., up to 10 days of stress deprivation) [36] which could explain the discrepancy in Mmp-13 response. Note, we did not examine Mmp-1 expression here, as it is not expressed in mice. Nevertheless, our findings indicate that F-actin destabilization corresponds to an increase in Mmp-3, a protease that plays a role in tendinosis [67]. The destabilization of F-actin is an important regulator of Mmps during disease processes like tendinosis.

We determined that tendon cells express several Tpm isoforms, including several known stress fiber-associated Tpms (Figure 3). Previously, Benjamin et al., (2002) indicated the presence of Tpm in tendon cells using Western blotting and immunofluorescence. However, Tpm antibodies are developed using peptides that correspond to specific exons that are highly conserved between various Tpms [63]. Thus, antibodies are specific to exons and not to particular isoforms. The particular (TM311) antibody used by Benjamin et al. recognizes an epitope on exon 1A of Tpm1 and 2 [63]. Therefore, it remained unclear which exact Tpm isoform was expressed in tendons. In our present study, we determined that tendon cells can express 9 Tpms: Tpm1.1, 1.6, 1.8, 2.1, 2.2, 3.1, 3.5, 3.12, 4.2. Based on our data, Benjamin et al. could have been probing for Tpms 1.1, 1.6, 2.1, 2.2. Tpm2.1 has been identified as a stress-fiber associated Tpm [46]. However, it is not present in all tendons, specifically tail tendons. Intriguingly, out of the nine Tpm isoforms expressed by tendon, we found that only three isoforms were consistently expressed in all tendons (tail, Achilles, plantaris) examined Tpm1.6, 3.1, and 4.2. This diversity of Tpm isoform may reflect the formation of specified F-actin networks in different tendon types to meet the unique tendon structures and mechanical functions [47]. Two of these isoforms, Tpm3.1 and 4.2, are known stress-fiber-associated Tpms. We chose to focus on Tpm3.1 as we previously showed it is a crucial regulator of F-actin stress fibers in lens epithelial cells [38]. The present study is the first, to our knowledge, to examine Tpm3.1 in tendon cells.

Tpm3.1 associates with F-actin in native tendon cells as well as *in vitro* cultured tendon cells to maintain F-actin stress fibers (Figure 3). The association of Tpm3.1 with stress fibers in tendon cells is in agreement with past studies on lens epithelial cells [38], human osteosarcoma (U2OS) [45, 46, 68, 69], human neuroblastoma (SK-N-BE(2) [70], and neuronal (B35) [62] cells. Tpm3.1 maintains F-actin networks through regulation of other actin associated molecules. It has been shown to promote the activation of non-muscle myosin activity [45, 46, 62, 70] as well as interfere with the association of severing molecules (i.e. cofilin) onto actin [62]. Here, we show that inhibition of Tpm3.1 results in the destabilization of F-actin stress fibers (Figure 5) leading to tendinosis-like molecule modulation (Figure 6 and 7). Surprisingly, as compared to stress deprivation of tendons, Tpm3.1 inhibition by TR100 treatment of cells had a more substantial effect on the genes tested. TR100 treatment not only decreased tenogenic expression but also enhanced chondrogenic genes, consistent with tendinosis [44]. Additionally, while stress deprivation modulated Mmp-3 but not Mmp-13, TR100 increased both Mmp-3 as well as Mmp-13. The greater response by TR100 treatment of cells could indicate the greater level of actin destabilization achieved by directly perturbing actin by TR100 treatment as compared to stress deprivation which may not have as vast an effect on F-actin network organization.

**Figure 6.**
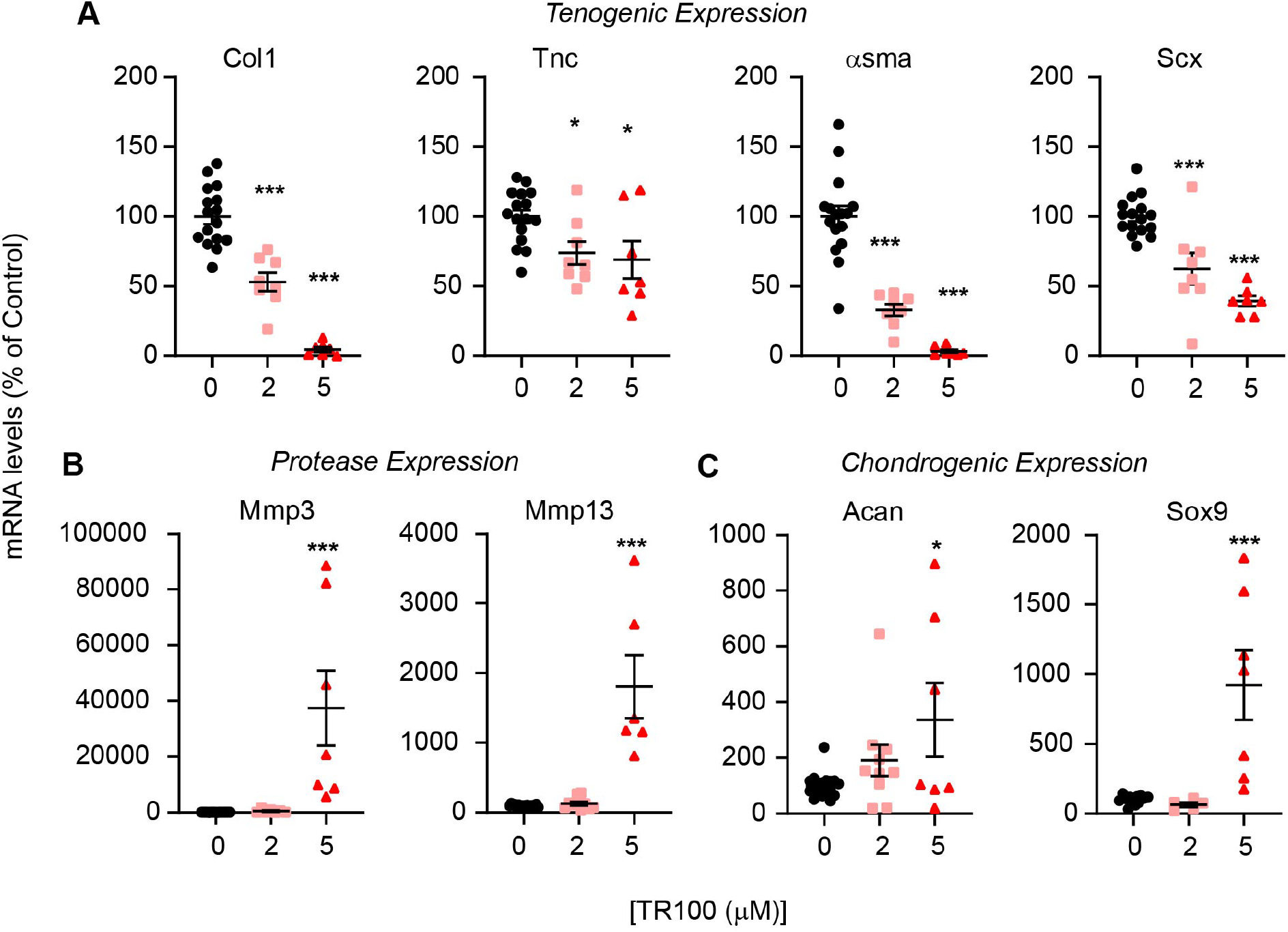
Destabilization of F-actin by inhibition of Tpm3.1 results in tendinosis-like gene modulation. Relative real-time PCR of tendon cells treated with 0, 2, or 5μM TR100 demonstrating (A) a reduction in tenocyte marker mRNA levels, and an (B) increase in protease mRNA levels, and (C) chondrogenic mRNA levels. Specific mRNA levels were normalized to Gapdh and normalized values are expressed as a percent of non-treated (0μM) controls. *, p < 0.05; ***, p < 0.001

**Figure 7.**
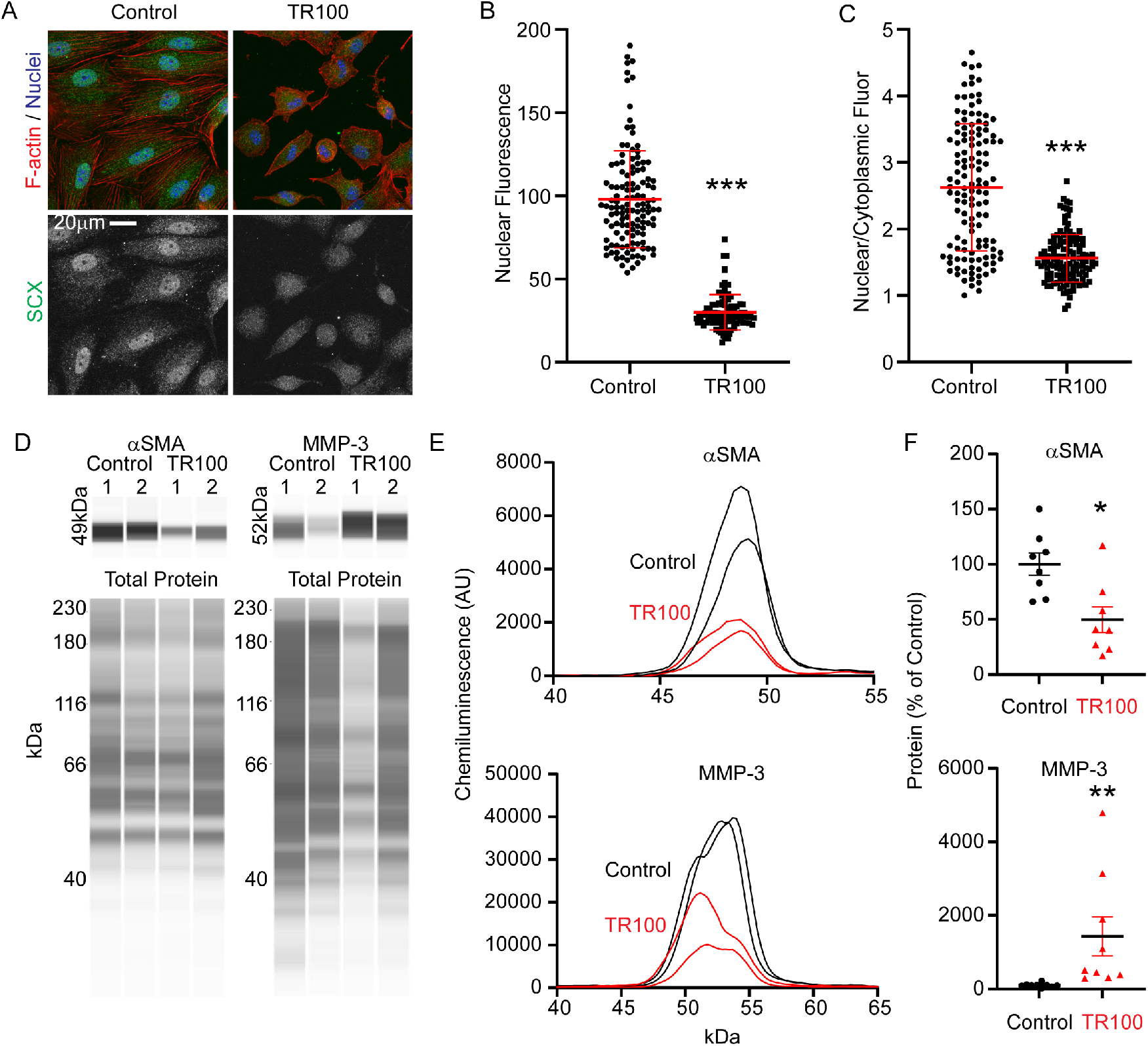
Modulation of select protein levels by TR100 (5μM) treatment. (A) Representative fluorescent confocal microscopy images for SCX. (B) Total nuclear fluorescence for SCX in individual cells, expressed as a percentage of mean nuclear fluorescence of control cells. (C) Calculated nuclear/cytoplasmic ratio of SCX for individual cells. Representative WES electrophoresis data showing (D) pseudo-western blots with corresponding (B) electropherograms for αSMA and MMP-3. (C) αSMA and MMP-3 protein after normalization to total protein. Protein levels were expressed as a percentage of untreated controls. *, p < 0.05; **, p < 0.01; ***, p < 0.001

It remains unclear how F-actin regulates gene expression in tendon cells. We have shown in chondrocytes and lens cells that F-actin depolymerization can directly regulate gene expression by controlling the cytoplasmic/nuclear shuttling of myocardin related transcription factor (MRTF) [38–40, 71]. The export of MRTF from the nucleus to the cytoplasm, caused by F-actin depolymerization, has been associated with downregulation of Col1, Tnc, and αsma expression as well as upregulation of Sox9 [38–40]. There is limited data that shows the regulation of Scx, Mmp-3, and Mmp-13 by MRTF. However, MRTF has been shown to regulate another Mmp, Mmp-12, in nucleus pulposus cells [72]. Intriguingly, in the present study, we identified a decrease in the nuclear proportion of transcription factor SCX by TR100 treatment. SCX is known to regulate sox9 expression, collagen synthesis, and the myofibroblast phenotype acquisition [60, 73–75]. Therefore, the decreased nuclear SCX may explain the upregulation of Sox9 as well as the downregulation of Col1, Tnc, and αsma mRNA levels that we found in the present study. To our knowledge, this is the first study to associate F-actin depolymerization regulation of SCX localization. Future studies are required to determine the mechanisms downstream of F-actin depolymerization that regulate gene expression.

Since we demonstrate that F-actin stability is regulated by Tpm3.1, this could lead to the speculation that the Tpm3.1 plays a role in tendinosis. Tendon overuse is often regarded as a cause of tendinosis. It has been suggested that tendon overuse causes a paradoxical stress deprivation of tendon cells [25]. In support of this mechanotransmission paradox, *ex vivo* overloading of rat tail tendon has been shown to result in cellular-matrix disruptions [26]. We speculate cellular-matrix disruption leads to cellular stress deprivation and destabilization of F-actin. We predict Tpm(s) to be involved in F-actin destabilization as Tpms have been shown to be mechanoregulated in tendon cells. Benjamin et al. have shown that *in vitro* mechanical stimulation of tendon cells elevate protein expression of Tpm [14]. Our future studies aim to delineate the role that Tpm3.1 plays in F-actin destabilization by stress deprivation using an *in vivo* model of tendinosis.

In conclusion, F-actin is a critical mediator of cellular phenotype and determining the regulation of F-actin networks by Tpms in tissues could lead to new therapeutic interventions that could prevent and/or reverse progression of disease in various tissue types including tendon.

## Acknowledgements

The authors thank Thomas Manzoni, Grace Emin, and Sofia Gonzalez-Nolde for critical reading of the manuscript. We also thank Sofia Gonzalez-Nolde for technical expertise in designing the real-time RT-PCR primers used in this study. Research reported in this project was supported by University of Delaware Research Fund – Strategic Initiatives (UDRF-SI) grant as well as the Delaware Center for Musculoskeletal Research from the National Institute of Health’s National Institute of General Medical Sciences under grant number P20GM139760.

